# MOJITOO: a fast and universal method for integration of multimodal single cell data

**DOI:** 10.1101/2022.01.19.476907

**Authors:** Mingbo Cheng, Zhijian Li, Ivan G. Costa

## Abstract

The advent of multi-modal single cell sequencing techniques have shed new light on molecular mechanisms by simultaneously inspecting transcriptomes, epigenomes and proteomes of the same cell. However, to date, the existing computational approaches for integration of multimodal single cell data are either computationally expensive, require the delineation of parameters or can only be applied to particular modalities.

We present a single cell multi-modal integration method, named MOJITOO (**M**ulti-m**O**dal **J**oint **I**ntegra**T**ion of c**O**mp**O**nents). MOJITOO uses canonical correlation analysis for a fast and parameter free detection of a shared representation of cells from multimodal single cell data. Moreover, estimated canonical components can be used for interpretation, i.e. association of modality specific molecular features with the latent space. We evaluate MOJITOO using bi- and tri-modal single cell data sets and show that MOJITOO outperforms existing methods regarding computational requirements, preservation of original latent spaces and clustering.

## 1 Introduction

The technological advances of high-throughput single cell sequencing enable us to characterize cellular heterogeneity of complex tissues for distinct molecular players of cells such as transcripts, proteins and chromatin^1^. The advent of multimodal technologies allow us to simultaneously measure two or more modalities at the same cells, i.e. RNA and open chromatin ^2–4^; RNA and protein^5^; and RNA, open chromatin and protein^6, 7^. These methods allow us to access how genetic information is associated at distinct molecular levels, i.e. the effect of DNA accessibility changes on gene expression or the expression of genes to proteins. However, data produced by each modality has quite distinct characteristics regarding their numerical values (e.g. low counts for open chromatin and variable count values for RNA and proteins levels), dimensionality (dozens for proteins, tens of thousands for genes, hundreds of thousands for open chromatin), and levels of data sparsity^8, 9^. These make the integrative analysis of multi-modal data a challenging task.

Here we are interested in the problem of estimating a shared latent space from parallel multiomic approaches, where two or more modalities are measured in the same cells. A few methods have been proposed for this problem. These follow two main frameworks: metric learning and latent variable learning. Weighted nearest neighbors (WNN) ^10^) and Schema^11^ explore, respectively, nearest neighbors and quadratic programming to estimate a single distance matrix representing the integrated multimodal data. Both approaches explore efficient algorithms, but do not explicitly provide models associating molecular features to the “latent space”. MOFA^12^, scAI^13^, totalVI^14^ and LIGER^15^ explore distinct methods for matrix factorization and estimation of shared latent spaces between modalities. Moreover, estimated matrices can be used for model interpretation, i.e., decomposed matrices can be used to associate molecular features with the latent space. Overall, these methods have a large number of free parameters including the size of the latent space (or rank of the low dimensional matrices). These methods require the optimization of the size of the latent space, which in turn increases computational costs. Moreover, the implementation of some methods (totalVI^14^ and scAI^13^) only allow integration of particular modalities (i.e., scRNA-seq and protein abundance for totalVI; scRNA-seq and scATAC-seq for scAI), while LIGER^15^ can only be used for two modalities and a subset of the molecular features need to be common in both modalities.

## 2 Approach

Here, we propose MOJITOO (**M**ulti-m**O**dal **J**oint **I**ntegra**T**ion of c**O**mp**O**nents), an efficient method that is based on canonical correlation analysis (CCA) to learn a shared latent space for any single-cell multimodal data protocol. The canonical components can be interpreted as factors and be used to characterize feature relevance by relating features across modalities (Fig.1). In contrast to matrix factorization methods, MOJITOO does not require the definition of parameters such as the rank of the matrix. Furthermore, it provides an approach to estimate the size of the latent space after a single execution of CCA. MOJITOO is provided as an R package and is compatible with common single cell pipelines for RNA, proteins (Seurat^10^ and ATAC modalities (Signac^16^).

**Figure 1.**
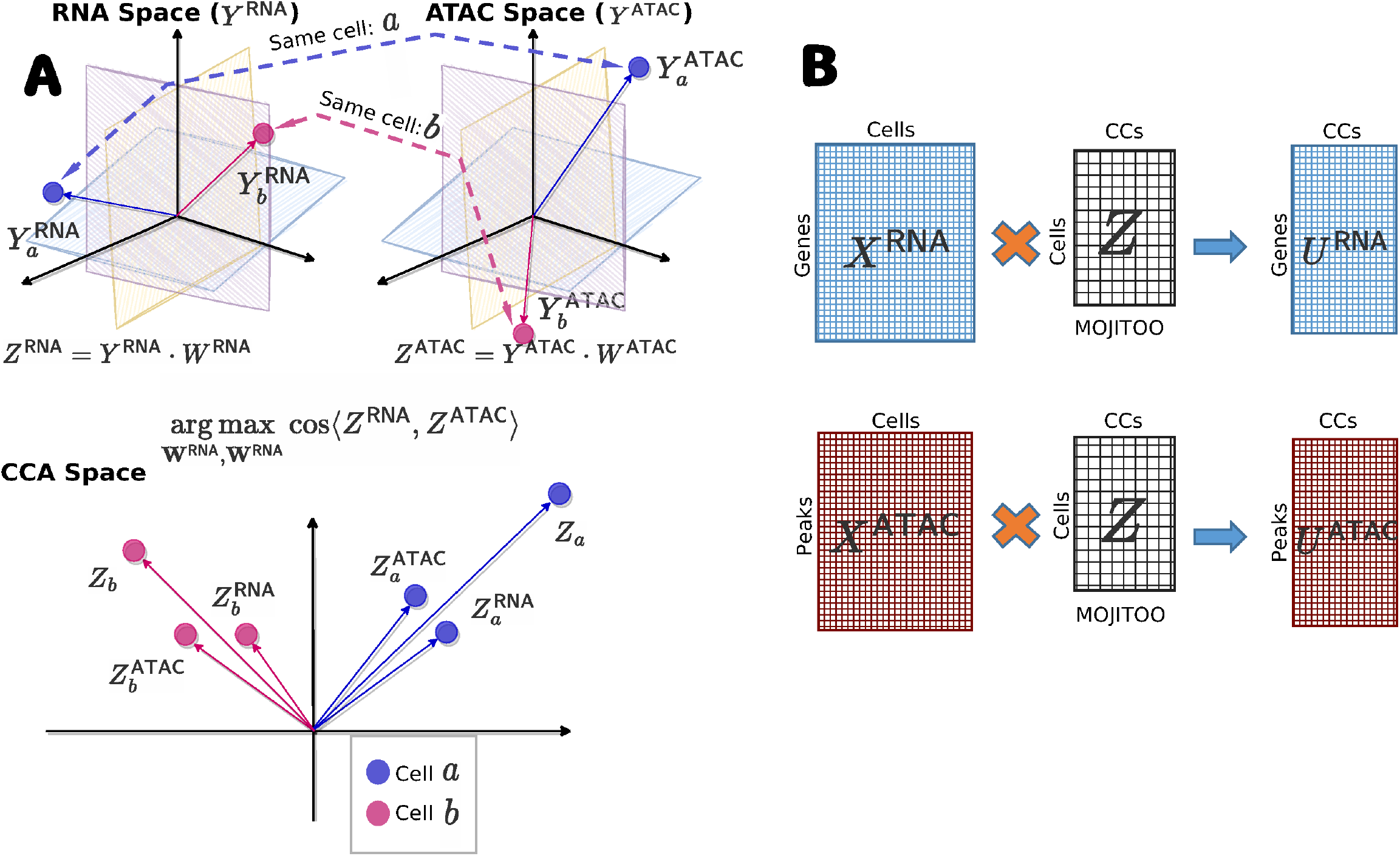
Schematic MOJITOO. **A**, MOJITOO receives as input two (or more) dimensional reduced matrices, where each matrix represents a particular molecular modality describing the same cells. In this example, we assume RNA and open chromatin (as measured by ATAC-seq) modalities are given. The main idea of MOJITOO is to use Canonical Correlation Analysis to find a set of canonical vectors *W*^ATAC^ and *W*^RNA^. Exploring a geometrical interpretation of CCA, MOJITOO finds canonical vectors such that the cosine similarity between latent dimensions in *Z*^RNA^ and *Z*^ATAC^ is maximized. A final representation *Z* can be obtained by adding the modality specific latent spaces. In the example, we show vectorial representations of two cells (*a* and *b*) in both original and latent spaces. **B**, An association between original features for each modality (*U*^RNA^ and *U*^ATAC^) can be obtained by multiplying original data representation per modality (*X*^RNA^ and *X*^ATAC^) with the shared latent space *Z*.

**Figure 2.**
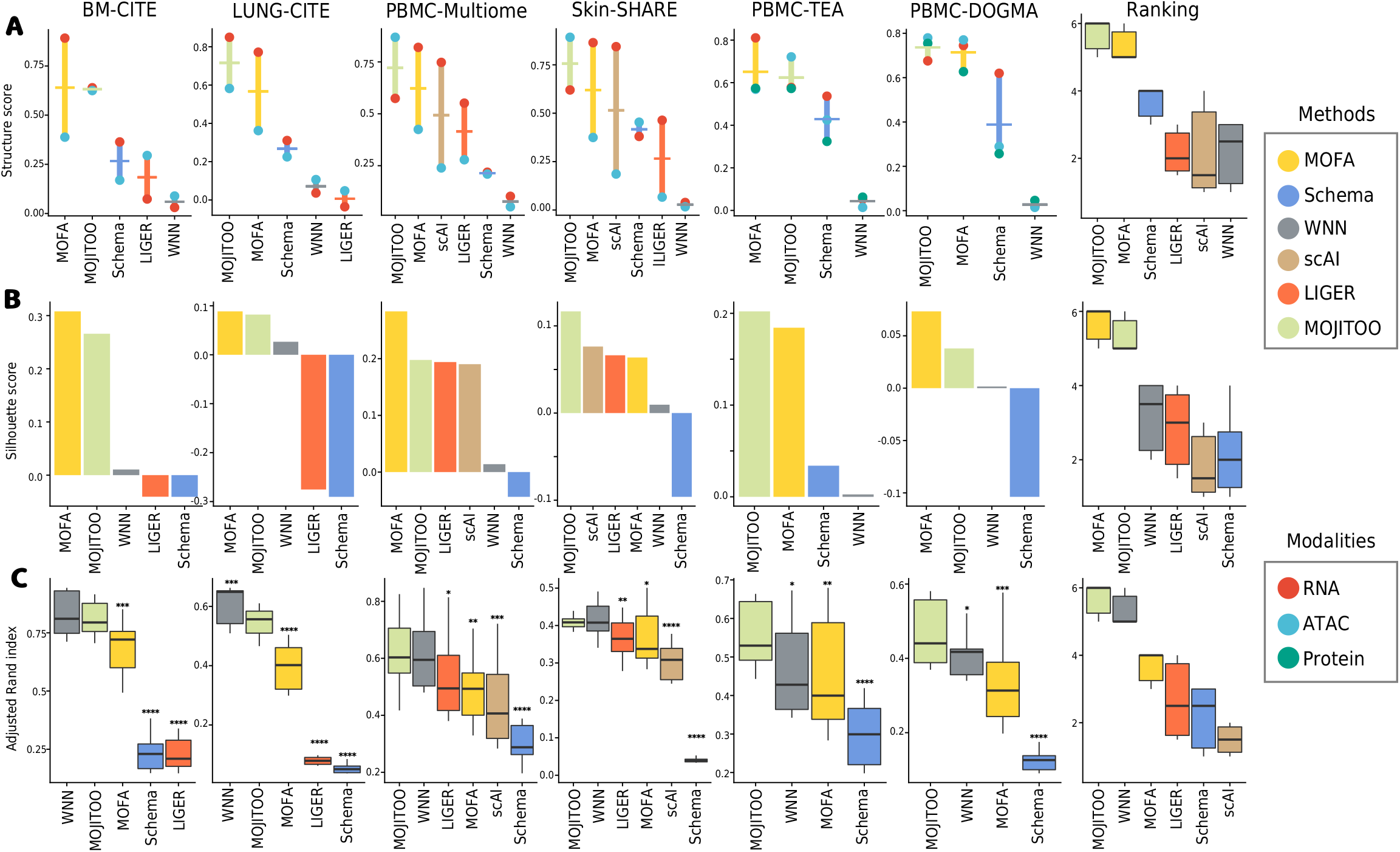
Benchmarking on data integration methods. **A**, We show the average (trace) and modality specific structure scores (dots) (*y*-axis) vs. methods (*x*-axis) for the six data sets. The last graph shows the combined ranking of the method over all data sets, where the highest rank indicates the best performer. **B**, Barplots showing silhouette score (*y*-axis) vs. methods (*x*-axis) for six benchmark data sets. The last plot shows the combined ranked per method. **C**, Boxplots showing ARI scores (y-axis) vs. methods (*x*-axis) for distinct clustering solutions for all six data-sets. Asterisks indicate *p*-values of <0.05(*), <0.01(**), <0.001(***), <0.0001(****) obtained via *t*-test comparing the ARI values of MOJITOO vs. other methods. The last boxplot shows the combined ranking for competing methods.

We evaluate MOJITOO and competing methods (WNN, MOFA, scAI, LIGER and Schema) in two bi-modal data sets with RNA and protein measurements ^17, 18^, two bi-modal data sets with RNA and ATAC-seq measurements^4^ and two tri-modal data sets with RNA, proteins and ATAC-seq measurements ^6, 19^ in regards to their ability to recover a shared space. The latent spaces are then evaluated with measures regarding the accuracy of clustering (adjusted Rand index), distance (silhouette score) and structure preservation, i.e. relation between shared space and original space of individual modalities^20^. Altogether, results show a superior performance of MOJITOO in both computational requirements and accuracy of estimated latent spaces. Moreover, we show how estimated canonical components can be used to interpret the underlying single cell data.

## 3 Methods

### 3.1 MOJITOO

MOJITOO takes as input a set of matrices from *m* modalities:

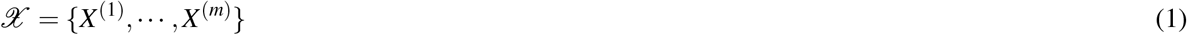

where 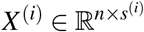 represents the data of a particular single cell modality, *n* represents the number of cells, and *s*^(*i*)^ represents the number of features in modality *i*. Here, we focus on multimodal data, where the cells are the same across matrices and there is no direct relation between the features of the distinct modalities.

#### 3.1.1 Reducing the dimension for each modality

We first obtain a dimension reduced matrix for each modality independently using a modality-specific approach:

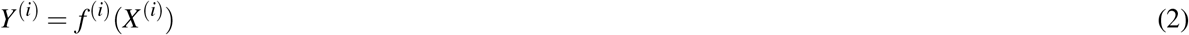

where 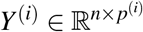 represents the low-dimensional matrix for modality *i, p*^(*i*)^ represents the number of dimensions and *f*^(*i*)^ represents the specific dimension reduction method for this modality. MOJITOO uses latent semantic indexing (LSI) for scATAC-seq and principal component analysis (PCA) for other modalities, as is usual in the literature^10, 16, 21^. The reason behind the use of dimension reduction is two fold. First, low-dimensional matrices reduce the computing time of the CCA analysis without impacting accuracy even when a small number of dimensions are used (30-50). Moreover, it allows to work directly on batch-corrected data, which is usually represented in a low-dimensional space^10, 22^.

#### 3.1.2 Learning a shared space with canonical correlation analysis with two modalities

MOJITOO aims to learn a shared latent space *Z* from the set of low dimensional matrices 𝒴 = {*Y*^(1)^, · · ·, *Y*^(*m*)^}

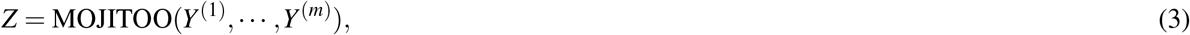

where *Z* ∈ ℝ^*n*×*k*^ represents the cells, *n* is the number of cells and *k* is the dimension of this latent space. When *Y* has two modalities, we first use CCA^1^ to project the matrices *Y*^(1)^ and *Y*^(2)^ to vectors 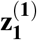 and 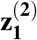:

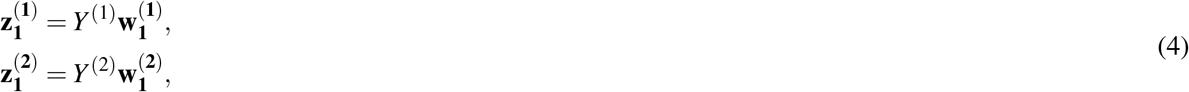

where 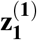 and 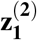 represent canonical components (CC). The vectors 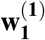 and 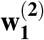 can be obtained by solving the following optimization problem:

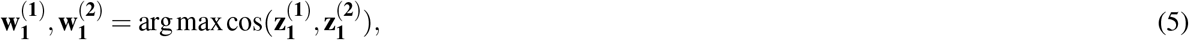

where 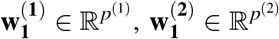 represent the first canonical weight vectors, and cos(·) is the cosine similarity between two vectors *a* and *b* defined by:

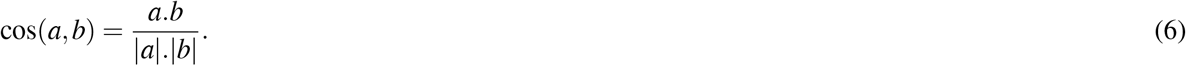

This is repeated 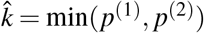 times, such that new canonical vectors are orthogonal to previously estimated vectors. These provide the matrices:

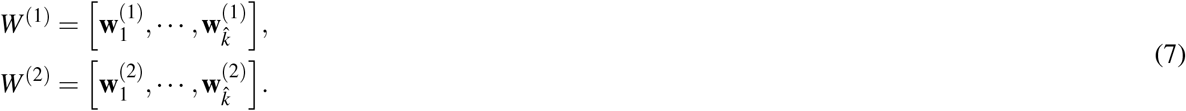

These can be used to estimate the modality transformed space as

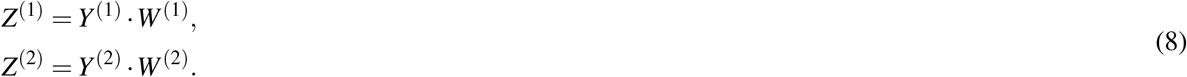

A unique latent space is obtained as

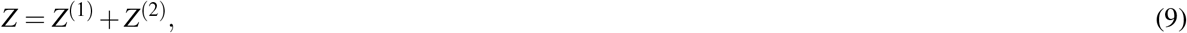

where *Z* ∈ ℝ^*n*×*k*^ and *k* is the number of canonical variables retained.

To further remove the noise from the latent space *Z*, we only keep highly correlated canonical components 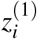 and 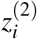 by measuring the Person correlation and using a student’s *t*-test for significance. The *p*-values are then corrected using BH(Benjamini Hochberg)^23^ and only canonical components with adjusted p-values < 0.05 are retained.

MOJITOO uses an algorithm based on generalized eigenvector decomposition^24^ to estimate the canonical components. MOJITOO has a time complexity of 𝒪(*max*{*p*^(1)^, *p*^(2)^}^2^ × *n*) for computing covariance matrices and 𝒪(min{*p*^(1)^, *p*^(2)^} × *p*^(1)^ × *p*^(2)^) for the eigenvector decomposition. As *n* (number of cells) is usually 100 times larger than *p*^(*i*)^ (number of reduced dimensions in *Y*^(*i*)^) the first term dominates the complexity.

Of note, CCA is one of the several steps in the integration algorithm of an earlier version of Seurat^25^. This had the objective to integrate distinct scRNA-seq experiments and CCA was performed in the common gene space, i.e. on transposed *Y*^(*i*)^ matrices and the objective was to find matching cells.

#### 3.1.3 Learning a shared space for multiple modalities

For the case that *Y* has more than two modalities, we perform the pairwise integration of modalities starting with the pair with highest dimensionality. The result of this CCA is then used for integration with the next modality. See algorithm 1 for a brief description, which receives a set of matrices {*Y*^(1)^, · · ·, *Y*^(*m*)^} with increasing dimensions *p*^(*i*)^ ≥ *p*^(*i*+1)^ as input. This heuristic algorithm was adopted to avoid the high computational costs of multiple CCA, which grows exponentially with the number of modalities.

##### Algorithm 1 Multimodal MOJITOO Algorithm

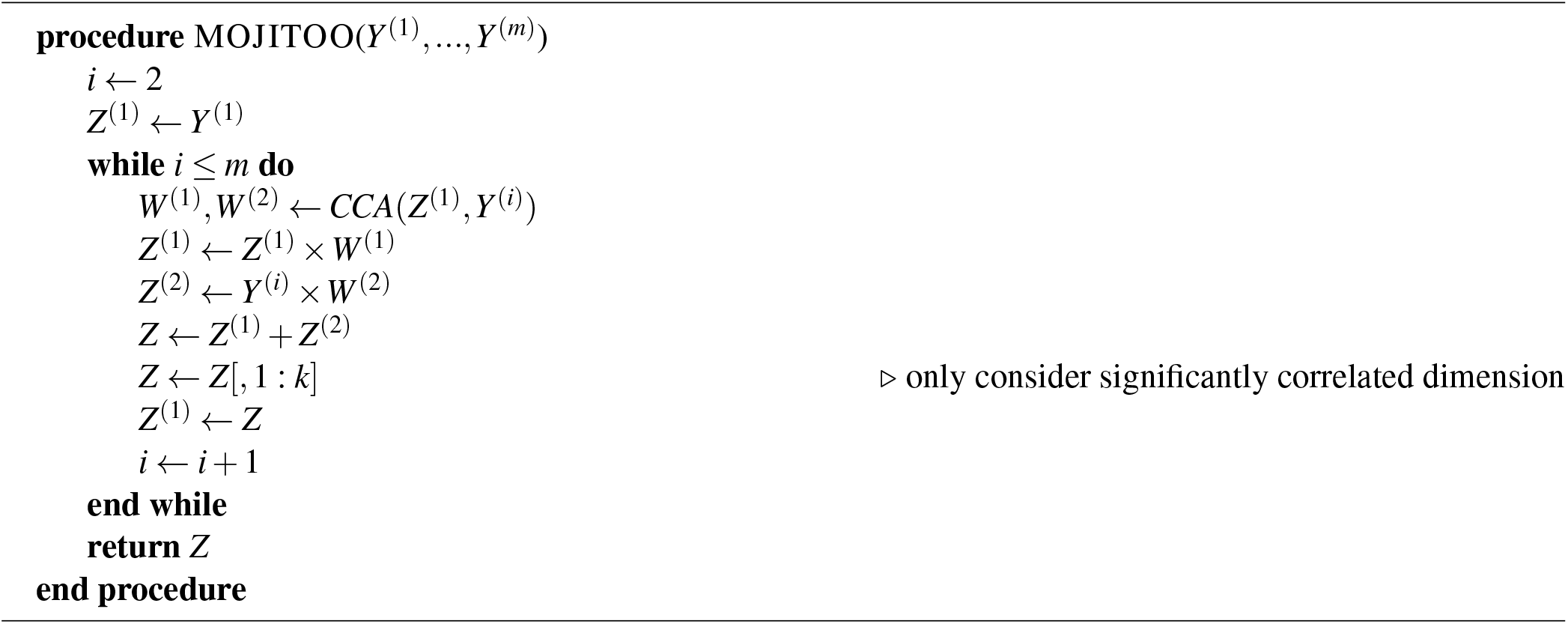

#### 3.1.4 Association of molecular features with latent space

We can use the estimated latent spaces to associate molecular features to the latent space *Z*. For example, let *X*^RNA^ ∈ ℝ^*n*×*s*^ be the gene expression matrix and *X*^ATAC^ ∈ ℝ^*n*×*t*^ be the peak matrix, where *n* is the number of cells, *s* is the number of genes and *t* is the number of peaks. We can obtain a feature associating molecular features to the latent space by

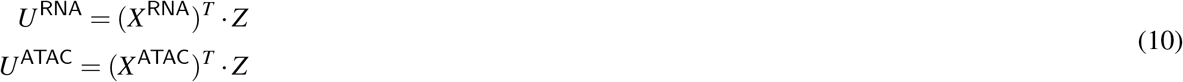

where *U*^RNA^ ∈ ℝ^*s*×*k*^ and *U*^ATAC^ ∈ ℝ^*t*×*k*^. The *i*th column of matrix *U*^RNA^ represent the scores of features in the *i*th canonical component.

### 3.2 Data sets

We make use of public multimodal data sets with two or tri-modalities in our evaluation. The first data set is single cell cite-seq data which measures single cell RNA and surface proteins simultaneously. The human bone marrow mononuclear cells (`BM-CITE`) data set contains full transcriptomes and 25 surface proteins for over 30,672 cells annotated in 27 cell types^17^. This data was obtained with the “LoadData(“bmcite”)” command from package SeuratData. Next, we applied the pre-processing pipeline. Another CITE-seq data used were the human peripheral blood mononuclear cells from lung (`LUNG-CITE`)^18^ with 52 surface proteins. It contains 10,470 cells annotated in 22 cell types. This data was obtained from here.

The next data set contains human peripheral blood mononuclear cells (`PBMC-multiome`) generated by the 10x multiome technology to measure gene expression (scRNA-seq) and chromatin accessibility (scATAC-seq) on the same cells. This data contains 11,787 cells with 13 cell types annotated by 10X Genomics. We use the scRNA-seq and scATAC-seq count matrices as provided by 10x genomics after processing with the cellranger pipeline obtained from the here. We also use a data set based on the SHARE-seq protocol measuring gene expression and chromatin accessibility of mouse skin cells (`SKIN-SHARE`)^4^. This data contains 34,774 cells, which are annotated as 23 cell types. We obtain the skin scRNA-seq and scATAC-seq counts and fragments files from the Gene Expression Omnibus under accession number (GSE140203).

A tri-modal data set of human PBMCs is measured with the DOGMA-seq protocol^6^. This provides RNA, ATAC and epitope sequencing of the same cells (`PBMC-DOGMA`). We use data under low-loss lysis condition, which contains 13,763 cells in 27 cell types. We download count matrices as provided by the authors [here](https://osf.io/6kr4v/). A second tri-modal dataset is based on human PBMCs measured with the TEA-seq protocol^7^. It contains transcripts, epitopes and chromatin accessibility of 25,517 PBMCs grouped into 12 cell types (`PBMC-TEA`). For this data set, we obtain original matrices and combine data from distinct wells from GEO (GSE158013). For scATAC-seq, we obtain an integrated matrix by combing peaks (allowing an extension of ±250bps). We finally intersect all barcodes from scRNA-seq, protein and scATAC-seq to obtain matrices in the same cell space. Characteristics of each of the six data sets are described in Table 1.

**Table 1.**
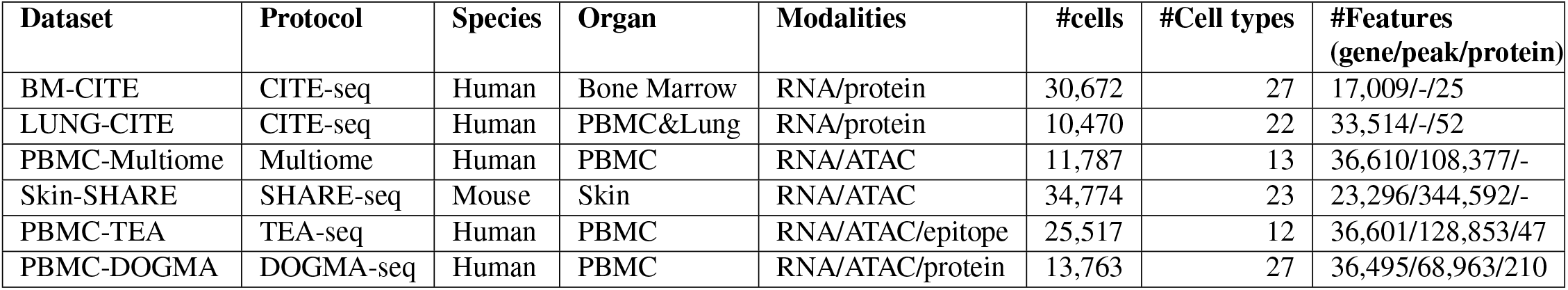
Major characteristics of multiomics data sets.

#### 3.2.1 Processing of single cell sequencing data

We perform a uniform pre-processing of all previously data sets starting from their count matrices. For scRNA-seq matrices, we adopt the standard Seurat 4 pipeline. First, we log normalize the data by calling the function NormalizeData with default parameters. Next we use FindVariableFeatures to find top 3000 variable features and run ScaleData. Finally, we use RunPCA to perform dimension reduction^10^ by keeping the first 50 PCs. For scATAC-seq, we adopt the standard pipeline from Signac^16^. We first run TF-IDF (term frequency - inverse document frequency) on the peaks. Next, we use RunSVD on the top features calculated by function FindTopFeatures with parameter min.cutoff=‘q0’, which provides an LSI dimension reduced matrix. We keep the first 50 dimensions, but we discard the first dimension as this is highly correlated to the number of fragments. For protein/epitopes, we adopt the standard Seurat 4 pipeline^10^. In short, we call NormalizeData with parameters normalization.method = ‘CLR’ and margin = 2 followed by ScaleData and RunPCA with 30 PCs using default parameters. For the PBMC-DOGMA data, we apply the harmony integration^22^ for RNA-seq and epitope data independently to integrate control and stimulated samples. For scATAC-seq, integration is performed by ignoring the first LSI dimension, which has a high correlation with the stimulation. We provide these input matrices to MOJITOO and WNN. For MOFA, we provide the normalized data, but without dimension reduction as in their tutorial (see below). Other competing methods provide their own functionalities for normalization and dimension reduction, which are used accordingly (see below). Time and memory requirements of pre-processing data are considered for the benchmarking of the respective method.

### 3.3 Benchmarking of integration methods

We use three distinct metrics to measure the accuracy of the methods. The `structure score` measures the similarity between two latent space structures^20^. It is based on the Pearson correlation of the pairwise Euclidean distance estimated on the shared (*Z*) and latent spaces (*Y*^(*i*)^) for each individual modality. This score indicates how well the shared space is related to the modality and the average values indicate how well integration worked. This metric is also employed by Schema^11^. We also evaluate the metrics concerning their distance representation using the `silhouette` score^26^. For this, we use the labels as provided by the cluster of the respective data set. We evaluate the use of Euclidean distance as ‘distance’ for the `silhouette` score. Finally, we evaluate the performance of methods regarding clustering. We perform Louvain clustering with varying resolution (parameter from 0.1 to 2.0) and estimate the adjusted Rand index (`ARI`) using cell labels^27^.

### 3.4 Execution of competing methods

#### MOFA

MOFA+^12^ uses Bayesian group factor analysis and variational inference to decompose individual modalities simultaneously by estimating a common latent factor matrix *Z*, as well as the weights for the transformation of the modalities to the latent space. MOFA+ includes a procedure to determine the optimal number of factors (dimension of the latent space) and has several hyper parameters for model regularization, detection of number of factors and learning rates. We execute MOFA with default parameters and followed their recommendations tutorial for the analysis of all data.

#### Schema

Schema^11^ explores metrics learning to re-weigh modality features through maximizing the agreement with other modalities. Specifically, it utilizes quadratic programming (QP) to learn a scaling transformation *u* for the primary matrix *X* such that pairwise distances of the transformation *u* * *x*_*i*_ (where * is coordinate-wise multiplication, for each *x*_*i*_ ∈ *X*) are highly correlated in other modalities. Schema has two main parameters: minimum desired correlation and number of random pairs. We run Schema using default parameters as in schema tutorial.

#### Seurat4 WNN

Weighted nearest neighbor (WNN)^10^ constructs single unified representation across multiple modalities. It first creates k-nearest neighbor (KNN) graphs for each modality based on the latent representation of each feature matrix. Next, it calculates affinities using the exponential kernel between a cell and the average NN for each modality. The latter is used to weigh cells. WNN has two major free parameters: the number of neighbors and scaling factor of the neighborhood kernel. We execute WNN, which is part of Seurat4, using default parameters. WNN does not provide a shared latent space, but we can use the weighted nearest neighbors graph to build a distance metric that can be used in all benchmarking evaluations.

#### scAI

scAI simultaneously decomposes transcriptomic and epigenomic data into multiple biologically relevant factors^28^. Its framework is similar to MOFA, but it can only cope with two modalities at a time. scAI uses a stability method to define the rank (size of the latent space) and has three main free parameters used for model regularization. We execute scAI in only bi-modal with RNA and ATAC-seq datasets with default parameters.

#### LIGER

LIGER^29^, which is based on non-negative matrix factorization, was originally proposed for data integration whenever modalities are in the same feature space. A newer variant of LIGER^15^ is able to perform integration, whenever there is some overlap between the features across modalities (shared features), i.e. protein and RNA expression of the same gene or gene accesibility scores for ATAC-seq. LIGER estimates a gene accessibility (ATAC-seq) matrix by counting the total number of ATAC-seq reads within the gene body and promoter regions(3kb upstream) for each gene per cell. An additional unshared feature matrix is further produced by binning the genome into bins of 100,000 bps and counting the overlap of these bins with peaks from the respective data set. LIGER has two major parameters: a regularization term and the number of factors (dimensions of the latent space). Regions associated to ENCODE Blacklist regions^30^ are removed. Moreover, LIGER can be only executed for bi-modal data sets.

## 4 Results

### 4.1 Benchmarking of multimodal integration methods

We evaluate MOJITTO and competing methods using six publicly available multimodal data sets with two or three modalities. These data sets have between 10,000 and 35,000 cells, 12 and 27 cell types and 25 to 344,492 features per modality (Table 1). We compare MOJITOO with MOFA^12^, WNN^10^, Schema^11^, scAI^13^ and LIGER^15^. Of note, some methods (scAI and LIGER) failed to be executed in some conditions, due to their inability to cope with more than 2 modalities or the lack of raw sequences for some of the evaluated data sets.

First, we evaluate algorithms regarding their structure preservation, i.e. the average similarity between the euclidean distances in the shared space and distances in the space of each modality^20^. Results indicate highest structure scores for MOJITOO (4 out of 6) followed by MOFA (2 out of 6). A ranking of the structure scores indicates MOJITOO as the best algorithm followed by MOFA and Schema (Fig. 3A). Interestingly, we observe that top competing methods (MOFA, Schema) tend to obtain higher structure scores for RNA and that MOJITOO has a structure score with lower variance across modalities. This suggests that the MOJITOO shared space captures information of all individual modalities more uniformly than MOFA and Schema, while MOFA and Schema have a tendency to focus on the RNA modality.

**Figure 3.**
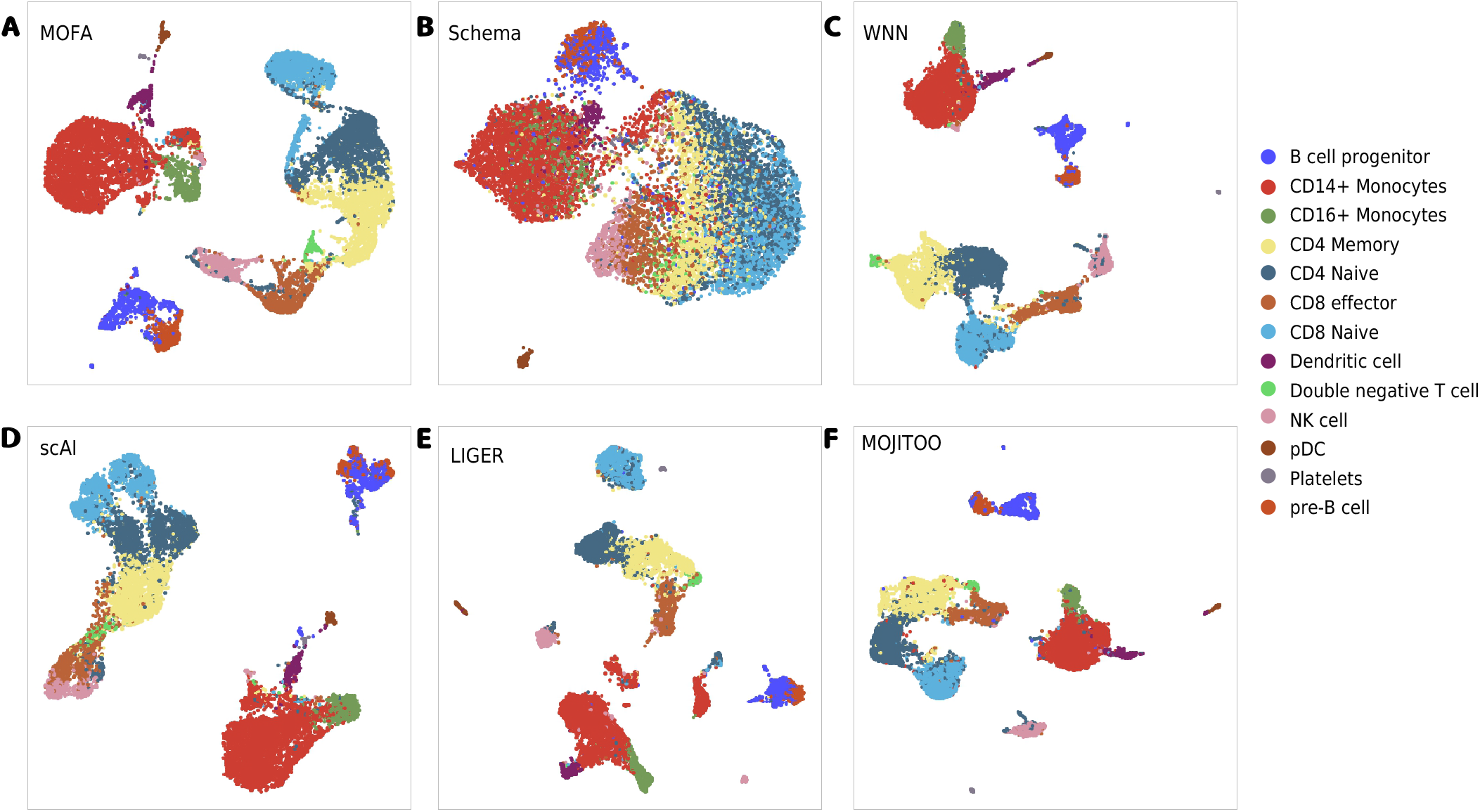
**A-F**, UMAPs showing cell type distribution derived from integration methods on PBMC-multiome dataset.

Next, we make use of the cell types reported in the original manuscripts introducing the single cell data sets as true labels for benchmarking. First, we use these labels to evaluate the silhouette scores by contrasting class labels with Euclidean distance matrices estimates on the shared space. Regarding silhouette, MOFA is best in 4 out of 6 data set, while MOJITTO is best in the other two data sets. MOJITOO obtains second rank in 4 out of 6 data sets and is ranked second in the overall ranking (Fig. 3B). Finally, we perform Louvain clustering at distinct resolutions (0.1 to 2.0) on the shared latent space. We then measure the agreement of clustering results with labels using the Adjusted Rand Index (ARI). Notably, MOJITOO obtains highest ARI in 4 data sets, while WNN is best in the two CITE-seq data sets (Fig 3C). MOJITOO has the highest overall rank followed by WNN. Examples of low dimensional embeddings obtained by distinct integration methods with the PBMC-Multiome data set are provided in Fig. 3.

A crucial aspect of single cell analysis is the computational resources needed for computation on an increasing number of cells. For this, we inspect the time and memory used in the largest data sets in our benchmark (SKIN-SHARE). To obtain curves, we down-sample the number of cells from 30,000 to 3,000 (Fig. 4A-B and Tables S1-S2). We observe that MOJITOO has the overall lowest computational requirement (2.4 minutes and 6.3 GBs) followed closely by WNN (3.74 minutes and 6.8 GBs). MOFA, on the other hand, required up to 67 minutes and 22.5 GBs for 30,000 cells, while scAI required 637 minutes and 75 GB of memory. These results reflect the fact that MOFA and scAI are based on complex matrix factorization algorithms, which require a computationally expensive optimization for the number of latent features. Altogether, results indicate MOJITOO has the best recovery of data structure and clustering results, while being the fastest and having the lowest memory footprint among all competing methods.

**Figure 4.**
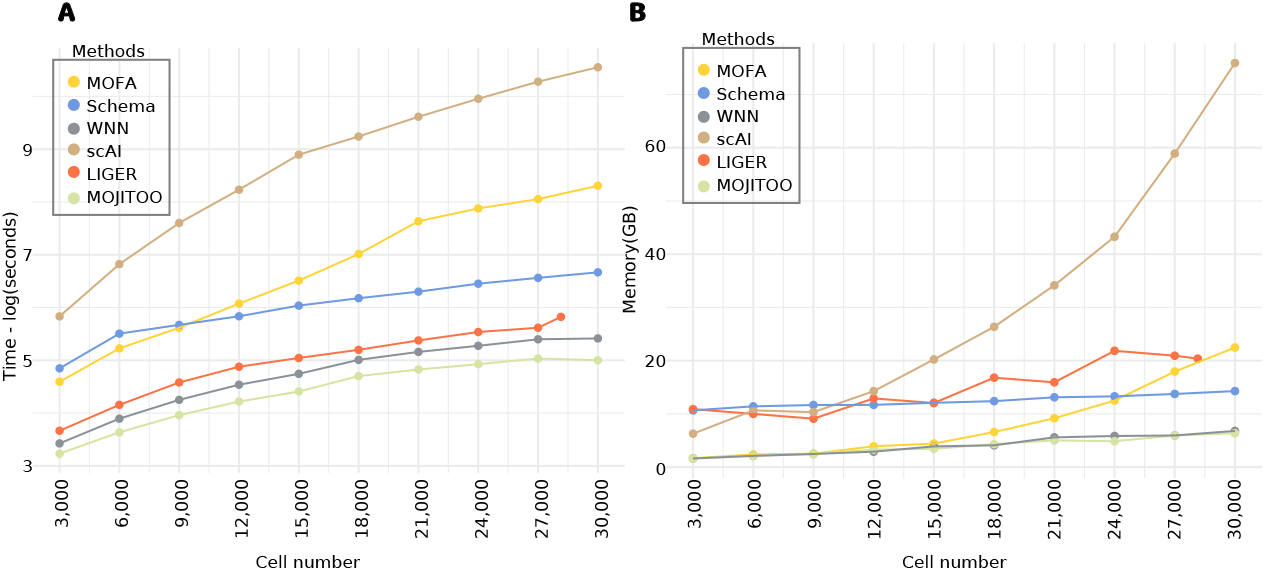
Time and Memory consumption on the Skin-SHARE. **A**, Line plots showing elapsed time (log of seconds) for each method (y-axis). **B**, Line plots showing peak memory (Gigabytes) required by each method (y-axis). In both **A-B**, the x-axis shows the number of cells used (randomly sampled) from the Skin-SHARE data.

### 4.2 Canonical vectors support the interpretation of multiome data

Additionally, we explore the use of the dimensions of the latent space (*Z*) as factors for interpreting the PBMC multiome data. We denote the latent features as canonical components (CC). As shown in Fig. 5, positive or negative values for the top CCs discern well all major cell types (Fig. 5). High values of CC1 are associated to myeloid cells (CD14+ and CD16+ monocytes and dendritic cells), while negative values are associated to T and NK cells (Fig. 5A). CC2 values discern B cell and plasmacytoid dendritic cells (pDC) from other cells, while CC3 differentiates B cells from pDCs (Fig. 5B-C). Further CCs capture subtle changes between major cell sub-types (Fig. 5D-E). CC4 and CC5 capture changes between naive T cells and active T CD8 and active T CD4 cells respectively, while CC5 captures differences between naive monocytes (CD14+) and activated monocytes (CD16+). Other smaller cell types (dendritic cells, platelets, double negative T cells and pre-B and progenitor B cells) can be characterized with further CCs (Figure S1).

**Figure 5.**
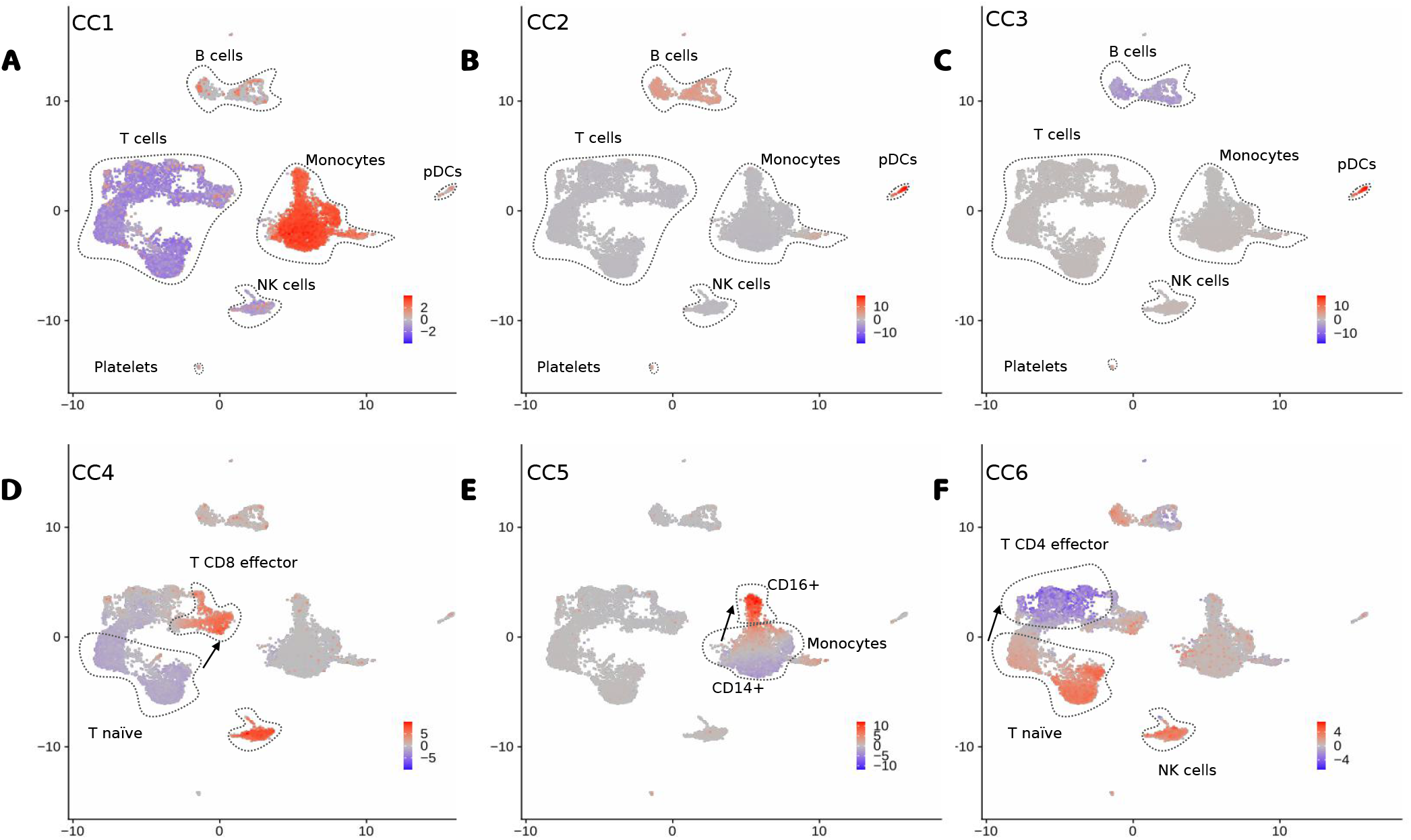
**A-F**, UMAP with the scores of CC1 to CC6. We highlight major cell types (or sub-types) associated to positive or negative CC scores and arrows indicate directions associated to the activation of particular immune cells.

Next, we explore the *U* matrices, which provide values associating molecular features with the latent dimensions (CCs). Indeed, the expression of genes with high CC1 values include monocyte genes as LYN and FCN1, while negative CC1 values are associated to T cell genes BCL11B and IL7R (Fig. 6A). Similarly, we observe that top ranked peaks with high or low CC1 scores have monocyte or T cell specific open chromatin. These include regions close to the T cell gene BCL11B (Fig. 6B). High CC2 value are associated with B cell genes IGHM and BCL11A, while low CC1 genes (BCBL11B and IL32) are associated with T cells (Fig. 6C). As before, we observe cell specific open chromatin patterns on top ranked ATAC-seq peaks associated with high and low CC2 values. Altogether, these results indicates that MOJITOO CCs can be used to capture major cell types of peripheral blood cells as well as to detect modality specific molecular features associated to these.

**Figure 6.**
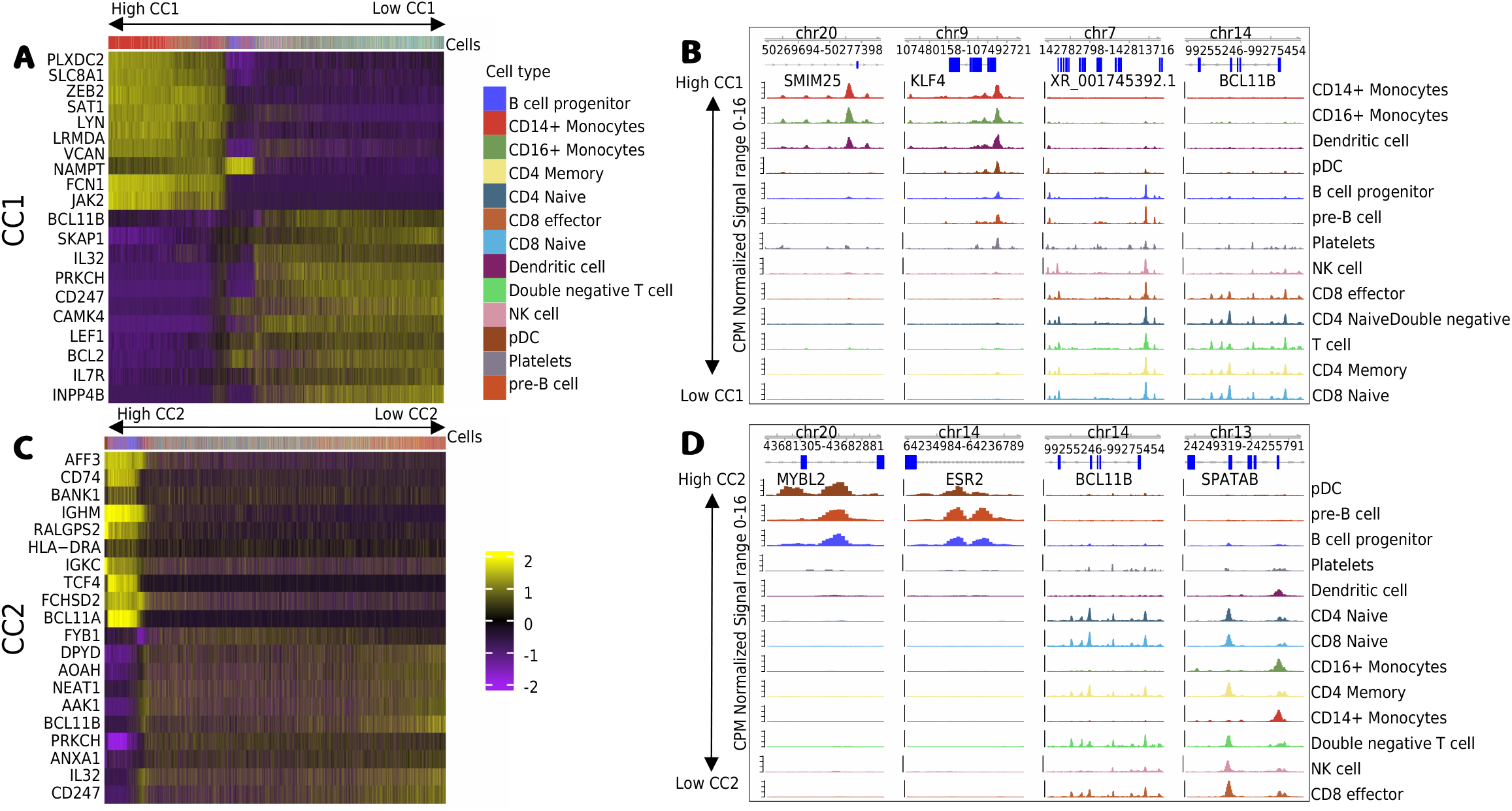
**A**, Heatmap with scores for the top 10 positive and negative genes for CC1 (*y*-axis) vs. cells (*x*-axis). Cells are ordered by CC1 scores (high to low). **B**, Genome browser tracks with top 2 positive and negative peaks for CC1. Tracks correspond to normalized cell specific pseudo bulk ATAC-seq profiles generated by deeptools^31^. Cell specific tracks are ordered by CC1 score (high to low). **C**, and **D**, show respectively the heatmap of top genes and the genome browser of top peaks for CC2.

## 5 Conclusion

We present here MOJITOO, which is a fast and parameter free method based on canonical correlation analysis for integration of multimodal single cell data of any protocol. A comprehensive analysis with six bi-modal and tri-modal multimodal data sets indicates that MOJITOO has the best performance regarding the preservation of the structures across modalities and the recovery of clusters, while it is ranked second regarding distance representation. Moreover, MOJITOO has the lowest time and memory requirements requiring 2.5 minutes and 6.4GB in the largest data set with 30.000 cells. WNN, which is the standard method for integration in Seurat, performed well on the clustering problem (2nd after MOJITOO) and had a low computational time, but had a poor performance in the structure preservation and silhouette scores. Moreover, WNN, which outputs a distance matrix on the shared space, does not provide latent features as MOJITOO or MOFA. MOFA performed well on the structure recovery and distance representation, but did not perform well on clustering and had one of the highest computational requirements being 20 times slower than MOJITOO and WNN and requiring 3.5 times more memory. The performance of MOFA reflects its model complexity, which includes the optimization of the size of the latent space. MOJITTO, on the other hand, explores the fact that CCA can be resolved within a single run of an eigen-decomposition and the choice of the final latent space can be performed as a posthoc step without the need of further model estimations.

Another interesting result is the fact the structure preservation scores are more uniform across modalities for MOJITOO than competing methods, while runner-up methods (MOFA and Schema) obtained highest scores for the RNA modality. This is possibly rooted on the analytical frameworks of these methods. CCA analysis explicitly finds canonical vectors with high correlation across modalities, while matrix factorization methods (MOFA and Schema) do not explicitly guarantee factors are uniformly well represented across modalities.

Finally, we highlight how a simple inspection of CCA derived latent spaces supports the biological interpretation and detection of relevant molecular features, as exemplified in the multiome PBMC data set. Future work includes further exploring the interpretability of MOJITOO, for example, by finding associations between molecular features across modalities as gene to peak links^21^. Another interesting topic is to investigate if differences in the modality specific space for given cell indicates biological properties of those. For example, in the Skin SHARE-seq data^4^, authors show that cells with changes in chromatin preceding changes in gene expression indicates cell differentiation.

## Code availability

Code and documentation are available on github: https://github.com/CostaLab/MOJITOO

## Acknowledgements

This project has been funded by the German Research Foundation (DFG) (project GE 2811/3-2) and the E:MED Consortia Fibromap funded by the German Ministry of Education and Science (BMBF).

## 6 Supplement

**Table S1.** Benchmarking experiments on SKIN-SHARE data set (time elapsed in minutes). Of note LIGER could only be executed with up to 28,147 cells.

| cells | LIGER | MOFA | MOJITOO | scAI | Schema | WNN |
| --- | --- | --- | --- | --- | --- | --- |
| 3,000 | 0.65 | 1.65 | 0.42 | 5.69 | 2.12 | 0.51 |
| 6,000 | 1.06 | 3.10 | 0.63 | 15.29 | 4.10 | 0.82 |
| 9,000 | 1.63 | 4.58 | 0.88 | 33.37 | 4.84 | 1.17 |
| 12,000 | 2.19 | 7.24 | 1.13 | 62.68 | 5.70 | 1.56 |
| 15,000 | 2.59 | 11.20 | 1.37 | 121.74 | 6.98 | 1.92 |
| 18,000 | 3.02 | 18.53 | 1.83 | 171.82 | 8.02 | 2.50 |
| 21,000 | 3.61 | 34.51 | 2.08 | 249.61 | 9.08 | 2.90 |
| 24,000 | 4.23 | 43.96 | 2.30 | 350.98 | 10.56 | 3.26 |
| 27,000 | 4.58 | 52.47 | 2.56 | 485.13 | 11.79 | 3.68 |
| 30,000 | - | 67.53 | 2.48 | 637.52 | 13.09 | 3.74 |

**Table S2.** Peak memory consumption in gigabytes. Of note LIGER could only be executed with up to 28,147 cells.

| cells | LIGER | MOFA | MOJITOO | scAI | Schema | WNN |
| --- | --- | --- | --- | --- | --- | --- |
| 3,000 | 10.88 | 1.66 | 1.61 | 6.27 | 10.66 | 1.61 |
| 6,000 | 9.99 | 2.37 | 2.11 | 10.65 | 11.41 | 2.11 |
| 9,000 | 9.09 | 2.51 | 2.46 | 10.32 | 11.66 | 2.46 |
| 12,000 | 12.90 | 3.90 | 3.28 | 14.26 | 11.68 | 2.89 |
| 15,000 | 12.04 | 4.42 | 3.43 | 20.22 | 12.07 | 3.88 |
| 18,000 | 16.80 | 6.58 | 4.31 | 26.36 | 12.39 | 4.09 |
| 21,000 | 15.92 | 9.17 | 5.01 | 34.14 | 13.11 | 5.58 |
| 24,000 | 21.85 | 12.49 | 4.85 | 43.27 | 13.29 | 5.84 |
| 27,000 | 20.93 | 17.94 | 5.94 | 58.91 | 13.74 | 5.93 |
| 30,000 | - | 22.47 | 6.34 | 75.92 | 14.28 | 6.79 |

**Figure S1.**
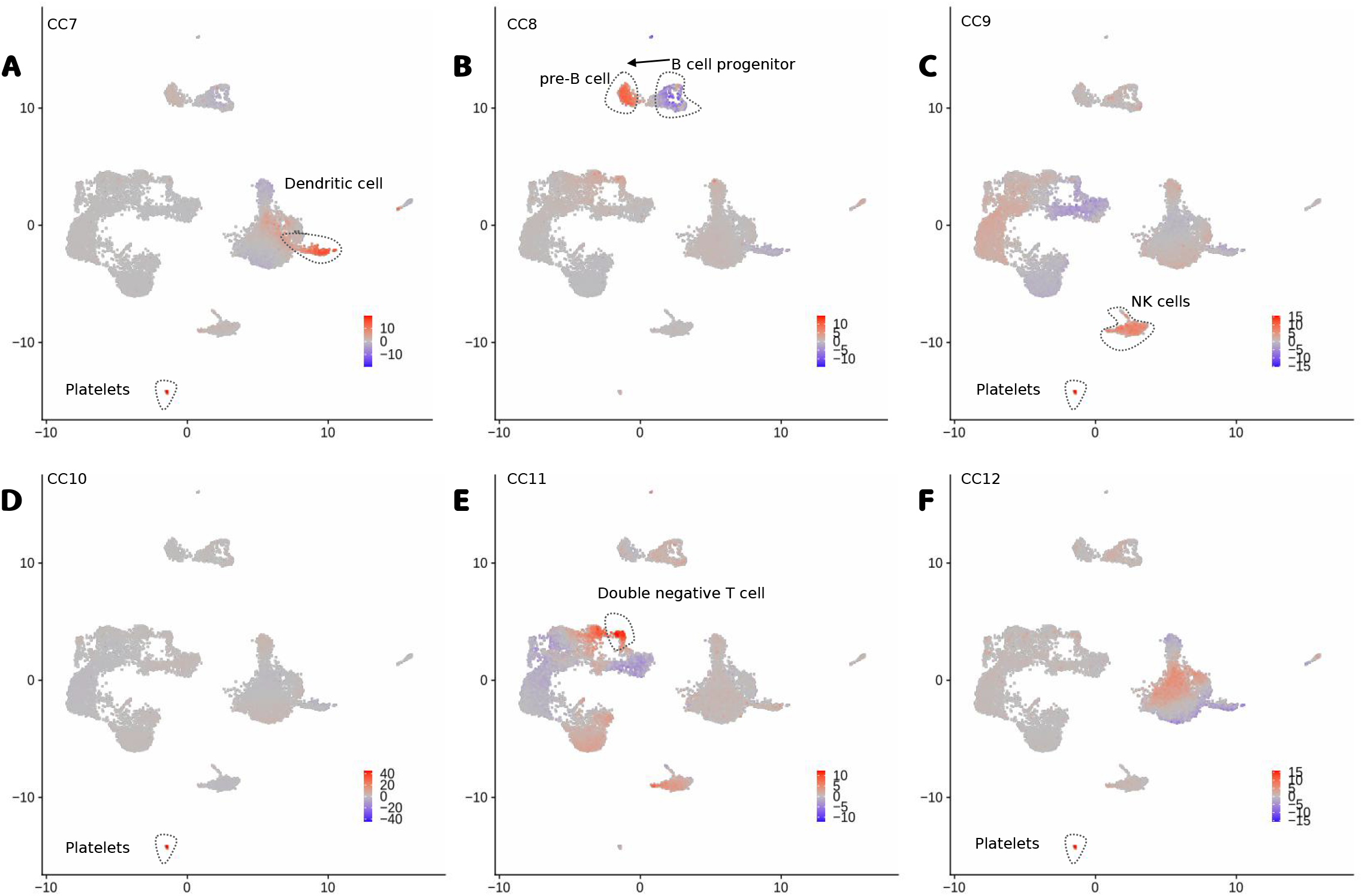
**A-F**, UMAP with the scores of CC7 to CC12. We highlight major cell types associated to positive or negative CC scores and the arrow represents a potential differentiation process.

## Footnotes

1 This notation is based on a geometrical interpretation of CCA.

